# Rapid olfactory discrimination learning in adult zebrafish

**DOI:** 10.1101/314849

**Authors:** Iori Namekawa, Nila R. Moenig, Rainer W. Friedrich

## Abstract

The zebrafish is a model organism to study olfactory information processing but efficient behavioral procedures to analyze olfactory discrimination and memory are lacking. We devised an automated odor discrimination task for adult zebrafish based on olfactory conditioning of feeding behavior. Presentation of a conditioned odor (CS+), but not a neutral odor (CS-), was followed by food delivery at a specific location. Fish developed differential behavioral responses to CS+ and CS- within a few trials even when odors were similar. The behavioral response to the CS+ was complex and included components reminiscent of food search such as increased swimming speed and water surface sampling. Appetitive behavior was therefore quantified by a composite score that combined measurements of multiple behavioral parameters. Discrimination behavior was robust in different strains and learned preferences could overcome innate odor preferences. These results confirm that zebrafish can rapidly learn to make fine odor discriminations. The procedure is efficient and provides novel opportunities to dissect the neural mechanisms underlying olfactory discrimination and memory.

## 1. Introduction

The zebrafish is an important vertebrate model to study neuronal circuit structure and function (Sumbre and de Polavieja 2014). Zebrafish larvae are small and relatively transparent, allowing for high-resolution optical measurements of neuronal activity throughout most of the brain. Even in juvenile and adult fish, activity patterns throughout major brain areas can be measured and manipulated by optical methods (Aoki et al. 2013; Fajardo et al. 2013; Jetti et al. 2014; Portugues et al. 2013; Rupprecht et al. 2016; Zhu et al. 2012). The small size of the zebrafish brain is also advantageous for dense reconstructions of neuronal connectivity (Friedrich et al. 2013; Wanner et al. 2016) and for genetic or chemical screens (MacRae and Peterson 2015). Zebrafish thus provide unique opportunities for quantitative analyses of brain function in wild type animals and in genetic disease models.

To understand the neural basis of behavior it is important to quantify behaviors of interest. At embryonic and early larval stages, zebrafish show primarily reflex-like sensory-motor behaviors with a limited potential for plasticity. Social behaviors and an increasing potential for associative learning emerge at later larval and juvenile stages (Buske and Gerlai 2011; Dreosti et al. 2015; Valente et al. 2012). Adult zebrafish and other teleosts show complex innate and cognitive behaviors (Abril-de-Abreu et al. 2015; Arganda et al. 2012; Brown et al. 2006; Buske and Gerlai 2011; Chou et al. 2016; Kalueff et al. 2013; Saverino and Gerlai 2008; Yabuki et al. 2016) including place preference and associative learning in different sensory modalities (Agetsuma et al. 2010; Aoki et al. 2013; Braubach et al. 2009; Darland and Dowling 2001; Doyle et al. 2017; Eddins et al. 2009; Lau et al. 2006; Mueller and Neuhauss 2012; Sison and Gerlai 2010; Xu et al. 2007). While various quantitative procedures have been established to analyze sensory-motor behaviors of zebrafish larvae there is an increasing demand for methods to study complex behaviors and learning in juvenile or adult zebrafish.

Adult zebrafish have been used to study principles of information processing in olfaction (Friedrich 2013). Imaging experiments demonstrated that odors evoke scattered but non-random spatial patterns of activity across the olfactory glomeruli, the input channels of the olfactory bulb (Friedrich and Korsching 1997). Neuronal circuits within the olfactory bulb decrease the overlap between activity patterns representing similar odors and stabilize odor representations against variations in stimulus intensity (Friedrich 2013; Friedrich and Laurent 2001; Niessing and Friedrich 2010; Zhu et al. 2013). Hence, processing of odor-evoked activity patterns in the olfactory bulb may support the classification of odor representations in higher brain areas. Further studies provide insights into transformations of spatio-temporal activity patterns between the output neurons of the olfactory bulb, the mitral cells, and telencephalic area Dp, the homolog of olfactory cortex (Blumhagen et al. 2011; Jacobson et al. 2018; Yaksi et al. 2009). To explore the impact of these computations on behavior, quantitative procedures are desired to analyze odor discrimination behavior and associative learning in adult fish. Ideally, such procedures should be automated and simple to implement in a standard fish facility.

Adult zebrafish can learn to associate different sensory stimuli with reward or punishment (Agetsuma et al. 2010; Aoki et al. 2013; Braubach et al. 2009; Darland and Dowling 2001; Doyle et al. 2017; Eddins et al. 2009; Lau et al. 2006; Mueller and Neuhauss 2012; Sison and Gerlai 2010; Xu et al. 2007). Two procedures have been described for olfactory conditioning of appetitive behaviors in adult zebrafish. Based on a procedure developed for catfish (Valentincic et al. 2000), Valentinčič and colleagues paired the infusion of an odor into a home tank with food delivery 90 s later (Miklavc and Valentinčič 2012). After approximately 30 training trials, the frequency of large-angle turns during the 90 s period after odor onset was significantly higher than the turn rate evoked by non-trained odors.Braubach and colleagues developed an appetitive olfactory conditioning procedure that follows a similar rationale (Braubach et al. 2009; Braubach et al. 2011). The results obtained using these approaches showed that zebrafish can learn olfactory associations and discriminations. However, both approaches require substantial technical resources that are not available in a standard zebrafish facility. Moreover, most experiments did not analyze odor discrimination. We therefore sought to develop a procedure for olfactory discrimination learning that is simple to implement and automate.

In order to minimize stress we chose an appetitive rather than an aversive conditioning paradigm. Odor discrimination tasks for rodents often involve unfamiliar behavioral components such as nose pokes or lever presses (Bodyak and Slotnick 1999; Frederick et al. 2009; Kay and Laurent 1999; Rinberg et al. 2006). We avoided such unfamiliar components and conditioned a familiar feeding behavior on an olfactory cue in an environment similar to the home tank. One odor stimulus (CS+) was followed by a food reward while another stimulus (CS-) had no consequence. A closely related strategy has recently been successful in auditory and visual conditioning of adult zebrafish without discrimination training (Doyle et al. 2017). Our behavioral paradigm includes the analysis of multiple behavioral components and resulted in rapid and robust olfactory discrimination learning, thus providing a basis for mechanistic analyses of olfactory processing and associative learning in zebrafish.

## 2. Methods

### 2.1. Animals

Zebrafish (Danio rerio) were raised and kept as groups in a standard facility at 26.5 – 27.5 °C on a 14/10h light/dark cycle. Fish were 7 – 10 months old and not selected for sex. Different wild-type strains and transgenic lines were used in separate cohorts. Experiments 1 and 3 were performed using wild-type fish (Abek × WIK). Experiments 2 and 4 were performed using fish that expressed halorhodopsin fused to YFP (eNpHR3.0YFP; (Gradinaru et al. 2010)) in GABAergic neurons. Experiment 5 was performed using a mixture of different wild-type and transgenic fish. All experimental protocols were approved by the Veterinary Department of the Kanton Basel-Stadt (Switzerland).

### 2.2. Experimental setup and odor application

Throughout the experiment, fish were kept individually in tanks that were usually custom-made from flat transparent polystyrole to avoid optical distortions by curved surfaces (typical dimensions: 29 × 9.5 × 7 cm; height of the watercolumn: ca. 6 cm). The tank was divided by a mesh into a front compartment containing the fish (~10 × 20 cm) and a rear compartment containing the suction tube for water removal (~9 × 10 cm). A feeding ring with a diameter of ~4.5 cm was made of silicon tubing and floated in a front corner (Fig. 1A). Food was delivered into the feeding ring from a remote location through a feeding tube. In most experiments food was pushed down the tube into the feeding ring by a custom device that applied computer-controlled pulses of pressurized air to the food delivery tube. In a subset of experiments, food was manually blown into the feeding ring using a plastic Pasteur pipette. Visual contact between the fish and the experimenter was avoided. Food was either a mixture of Gemma Micro 300 (Skretting) and crushed food flakes (Tetramin, Tetra) or powder food (SDS100). Food was delivered in small portions that were usually consumed within <30 s.

**Fig. 1.**
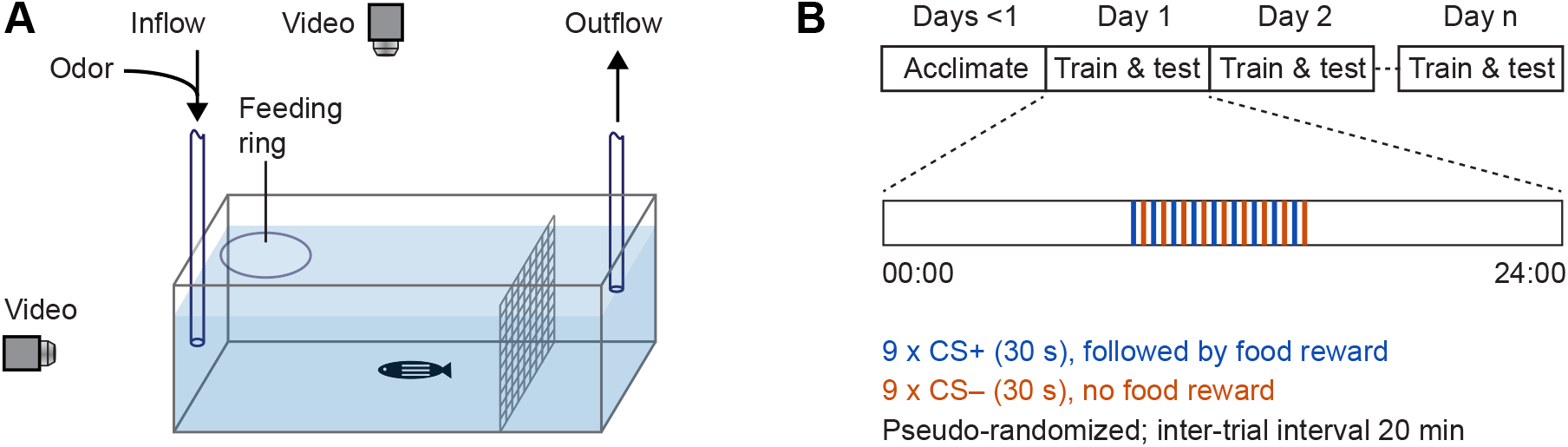
Experimental setup and protocol. (A) Schematic of experimental setup. (B) Experimental schedule.

Tanks were perfused with water from the fish facility using a peristaltic pump at a flow rate of 25 ml/min. The flow was constantly on for approximately 7 h on each day during the period when odor and food stimuli were delivered. Outside this time window and during the acclimation period the flow was off. Water entered the tank through an inflow tube (inner diameter, 1.6 mm) in the front at approximately at half-height in the water column. Water was removed from the tank by suction through an outflow tube at the rear of the tank and discarded. To minimize fluctuations of the water level, a pipette tip (200 μl) was attached to the tip of suction tube and positioned at the desired height.

Olfactory stimuli were amino acids, which are natural odorants for aquatic animals (Carr 1988). Odors were prepared freshly on each day by diluting a stock solution (6 × 10^-3^ M) in fish water to a concentration of 6 × 10^-5^ M. Odors were delivered into the tank by switching the perfusion for 30 s from a reservoir containing fish water to a reservoir containing odor solution. Switching was achieved using computer-controlled, motorized valves (WTA-2K-3MFE-3, Takasago Electric, Inc.) and valve controllers (<u>ValveLink8.2^®^ Controller, AutoMate Scientific</u>). Video analyses using a dye showed that the stimulus distribution during the first 30 s of application was inhomogeneous and discontinuous with highest concentrations near the inflow tube. Subsequently, the stimulus distribution equilibrated throughout the tank. The nominal odor concentration assuming even dilution in the tank was ~4.5 × 10^-8^ M, which is at or below the detection threshold of adult zebrafish for amino acids (Michel and Lubomudrov 1995; Miklavc and Valentinčič 2012). Hence, stimulus concentration was reduced to levels below threshold by dilution and water exchange (Miklavc and Valentinčič 2012).

Experiments were performed in up to four tanks in parallel, each containing a single fish. Tanks were separated by opaque screens, illuminated from below by infrared light, and filmed simultaneously by two orthogonal cameras (Fig. 1A). Sufficient video quality was achieved using standard web cameras at video rate (30 frames per second). Simple devices such as tennis balls or shock absorbers were used to partially isolate the tabletop from vibrations. The setup was placed inside an opaque enclosure which was lined with sound-absorbing foam in most experiments.

### 2.3. Experimental schedule

Fish remained in the experimental tanks after transfer from the facility for the duration of the experiment. During an initial acclimatization period of 1 – 3 days, fish received neither food nor odor stimuli. When training started, odors were applied at inter-trial intervals (ITIs) of 20 min. One odor (conditioned stimulus, CS+) was followed by food application (unconditioned stimulus, US) 30 s after stimulus onset while a second odor (CS-) was not followed by food application. Usually, a total of nine CS+ and nine CS- were delivered per day in alternating fashion (Fig. 1B). Hence, training extended over a period of six hours per day, usually starting at approximately 09:00 h. Fish were trained for a total of 2 – 9 days. During training, behavior was quantified in each trial during the 30 s period between odor onset and food delivery. Four fish of the same strain and age were usually trained in parallel on the same two odors A and B. Training was balanced such that two fish received odor A as CS+ and B as CS- while assignments were reversed for the other two fish.

### 2.4. Quantitative analysis of behavior

Orthogonal video recordings were analyzed to track the 3D position of each fish as a function of time (Fig. 2A). In each frame, the fish was segmented and represented by a point at the center of gravity. The following behavioral parameters were then extracted from the 3D trajectories: (1) Swimming speed. Instantaneous swimming speed was calculated as the displacement of fish between successive video frames. (2) Relative z-position. This parameter quantifies the relative position in the water column along the z-direction. 0 represents the bottom of the tank; 1 represents the water surface. (3) Residence in the reward zone. This binary parameter is 1 when the fish is located in the reward zone and 0 otherwise. The reward zone is defined as a rectangular area bounding the feeding ring and spanning the shorter axis of the experiment tank, covering approximately one third of the tank’s footprint. (4) Surface sampling events. A surface sampling event is defined as the crossing of a threshold in the z dimension below the center of the feeding ring. The threshold was set at approximately 70% of the height of the water column. Surface sampling events are usually discrete and occur when fish collect food from the water surface but are rare otherwise. (5) Distance to inflow. This is the 3D distance between the fish and the opening of the inflow tube. (6) Circling. This behavior refers to stereotyped circular swimming along the walls of the tank. We observed that such a swimming pattern is sometimes maintained for extended periods of time when fish are undisturbed but the behavior is interrupted by salient sensory input. To quantify circling we computed power spectra from measurements of the position along the long axis of the tank using a 30 s Tukey window and quantified the relative power in a low-frequency regime that corresponded to the frequency of circular swimming (0.029 – 0.146 Hz) in one-second time bins. Python-based software for automated video analysis is available at https://github.com/i-namekawa/TopSideMonitor.

**Fig. 2.**
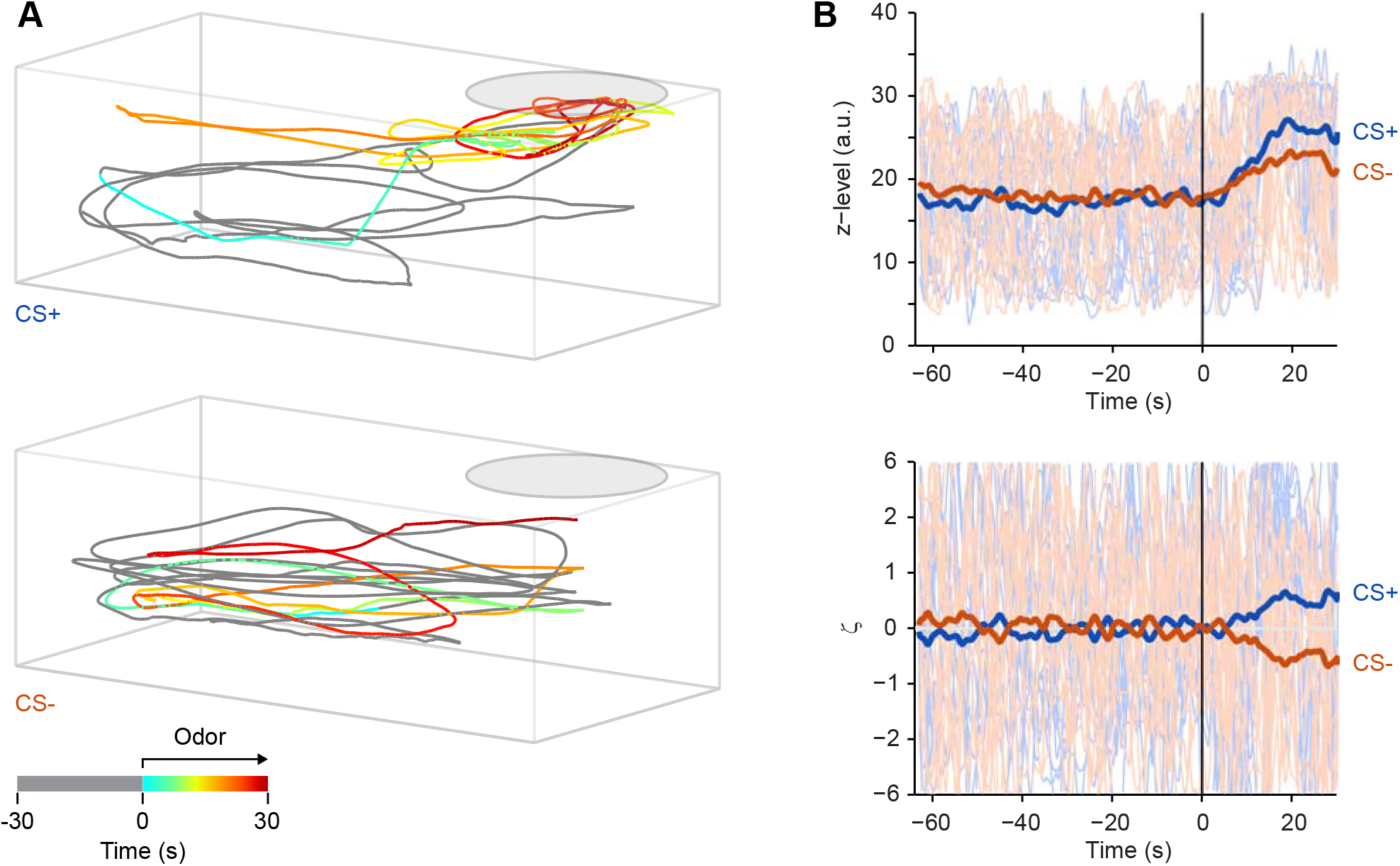
Analysis of behavior. (A) Examples of swimming trajectories prior to and during presentation of CS+ (top) and CS- (bottom) at the end of training. Each trajectory is 60 s long, comprising 30 s before and 30 s after odor onset, prior to food delivery. Time is color coded; pre-odor time is gray. (B) Top: time series showing one behavioral variable (z-level) as a function of time for all trials in an experiment; thick lines show averages. Odor was delivered at t = 0. Bottom: Same time series after transformation to ζ scores.

Measurements of each behavioral parameter in each video frame yielded one time series of values per parameter and trial. We defined three time windows for analysis: (1) *Baseline time window*. This window comprised the 30 s prior to odor onset and served as a baseline in each trial. (2) *Response time window*. This window was defined as the time between odor onset and the time of food delivery in CS+ trials or the equivalent time in CS- trials. This time window was usually 30 s long and used to assess behavioral responses to odor stimulation. (3) *Reference time window*. This time window was defined as a time window immediately prior to the baseline window with a length equal to the response window. This window served to assess trial-to-trial variability of behavior in the absence of stimulation. Note that the response window and the reference window were equidistant in time from the baseline window.

Because spontaneous behavior can fluctuate between trials we quantified the change in parameter values relative to a baseline in each trial. The baseline was defined as the mean value of a parameter during the baseline time window and subtracted from the time series. Time series were then averaged over all CS+ and over all CS- trials and the mean of these averages was subtracted from each individual time series. To quantify trial-to-trial variability in the absence of stimulation the standard deviation of the time-averaged values during the reference window across trials was calculated. The mean-subtracted time series were then normalized to this variability measure for each fish. As a consequence, the family of time series for each behavioral parameter was centered on zero and normalized by a measure of pre-stimulus variability (Fig. 2B). We refer to these transformed time series values as ζ scores because they are closely related to z-scores: positive values represent behavioral responses larger than the mean response to all stimuli, negative values represent behavioral responses smaller than the mean, and the absolute value reflects response magnitude relative to a variability measure. ζ scores differ from z-scores because they are normalized to the variability during a fixed pre-stimulus time window. As a consequence, ζ scores reflect the change in behavior relative to a baseline but they cannot be interpreted quantitatively as a z-score.

In order to obtain a single measure per trial, ζ scores were time-averaged over the response time window. The sequence of these averaged ζ scores over successive trials is referred to as a trial series of ζ scores.

## 3. Results

### 3.1. Olfactory discrimination learning: method and basic observations

We developed an experimental paradigm to establish associations between odors and food. Individual zebrafish were kept in tanks with continuous water perfusion and a feeding ring floating in one corner (Fig. 1A). Every 20 min, one of two amino acid odors (CS+, CS-; 6 × 10^-5^ M) was introduced into the perfusion for 30 s (Fig. 1B) and slowly removed thereafter by dilution and water exchange (Methods). Thirty seconds after the onset of the CS+, a few small food pellets were delivered into the feeding ring while the CS- was not rewarded. Usually, nine trials with each stimulus were alternated on each day. We hypothesized that fish learn to respond to the CS+, but not to the CS-, with anticipatory food search behavior.

Fish were tracked in 3D using two orthogonal video cameras and custom software. In trained fish, swimming trajectories showed obvious differences after delivery of the CS+ or CS- (Fig. 2A). Upon presentation of the CS+, fish often increased their swimming speed and spent more time near the feeding ring, consistent with previous observations in other olfactory conditioning paradigms (Braubach et al. 2009; Miklavc and Valentinčič 2012). In addition, fish often elevated their position in the water column, sampled the water surface as during feeding, and sometimes approached the inflow tube. Stereotyped cyclic swimming along the wall of the experiment tank, which usually occurred during inter-stimulus intervals, was often interrupted. These behavioral changes were less pronounced or absent upon presentation of the CS-.

To quantify these observations we analyzed parameters of the 3D swimming trajectory that reflect different behavioral components. Analysis tools were developed to automatically quantify (1) the instantaneous swimming speed (“Speed”), (2) the relative height of the fish in the water column (“z-level”), (3) the distance of the fish to the water inflow (“Distance”), (4) the frequency of surface sampling events (“Surface”), (5) the probability that the fish is located in a rectangular area around the feeding ring (“Area”), and (6) the amount of stereotyped, cyclic swimming (“Circling”). For each parameter and trial, time series of measurements were obtained. Measured values were transformed by (1) subtraction of the mean value during a 30 s “baseline time window” immediately before stimulus onset, (2) subtraction of the time series averaged over all trials, and (3) normalization to the inter-trial variability in a “reference time window” immediately prior to the baseline time window (Methods). This transformation results in a dimensionless measure that is normalized to pre-stimulus variability and centered on the mean at each time point (Fig. 2B). We therefore refer to these measures as ζ scores because they are closely related to z-scores (Methods). The signs of ζ scores for “Distance” and “Cycling” were inverted so that positive (negative) ζ values always reflected responses that were more (less) appetitive than the mean for all parameters. In order to analyze learning curves over trials, ζ scores were time-averaged during the 30 s between odor onset and food delivery (“response time window”) to obtain one value per trial. Medians were then calculated over the trials of each day to obtain one value per day.

### 3.2. Experiment 1: extended discrimination training

We initially trained 12 fish using Ala and Trp as odor stimuli (experiment 1; 9 CS+ trials and 9 CS- trials per day). Fish were trained for a total of nine days (81 trials for each stimulus). Assignments of odors as CS+ or CS- were balanced over fish. ζ scores of all parameters were significantly different between CS+ and CS- trials (Fig. 3A). Hence, fish learned to discriminate between the CS+ and the CS-.

Visual observations of behaving fish further suggested that the contribution of different components to the overall behavioral response varied substantially between trials and individuals. For example, a fish may respond to the CS+ primarily with fast swimming in one trial but with surface sampling in the next trial. Consistent with this observation, trial-by-trial correlations between most parameters were relatively low (Fig. 3B; mean correlation coefficient ± s.d.: 0.00 ± 0.27). Hence, combining multiple behavioral measurements may improve the detection of learned behavior because different parameters convey non-redundant information. Alternatively, combining measurements may increase noise. To explore these possibilities we computed a composite measure of behavior, ζ_comp_, based on Stouffer’s method for the combination of z-scores (Stouffer et al. 1949). As in Stouffer’s method, ζ_comp_ is the sum over all ζ scores normalized by the square root of the number of ζ scores (n = 6). This procedure enhanced, rather than decreased, the statistical significance (Fig. 4A). Hence, the combined analysis of multiple behavioral parameters enhanced the detection of learning-induced changes in behavior, presumably because the conditioned behavior comprises multiple components.

**Fig. 3.**
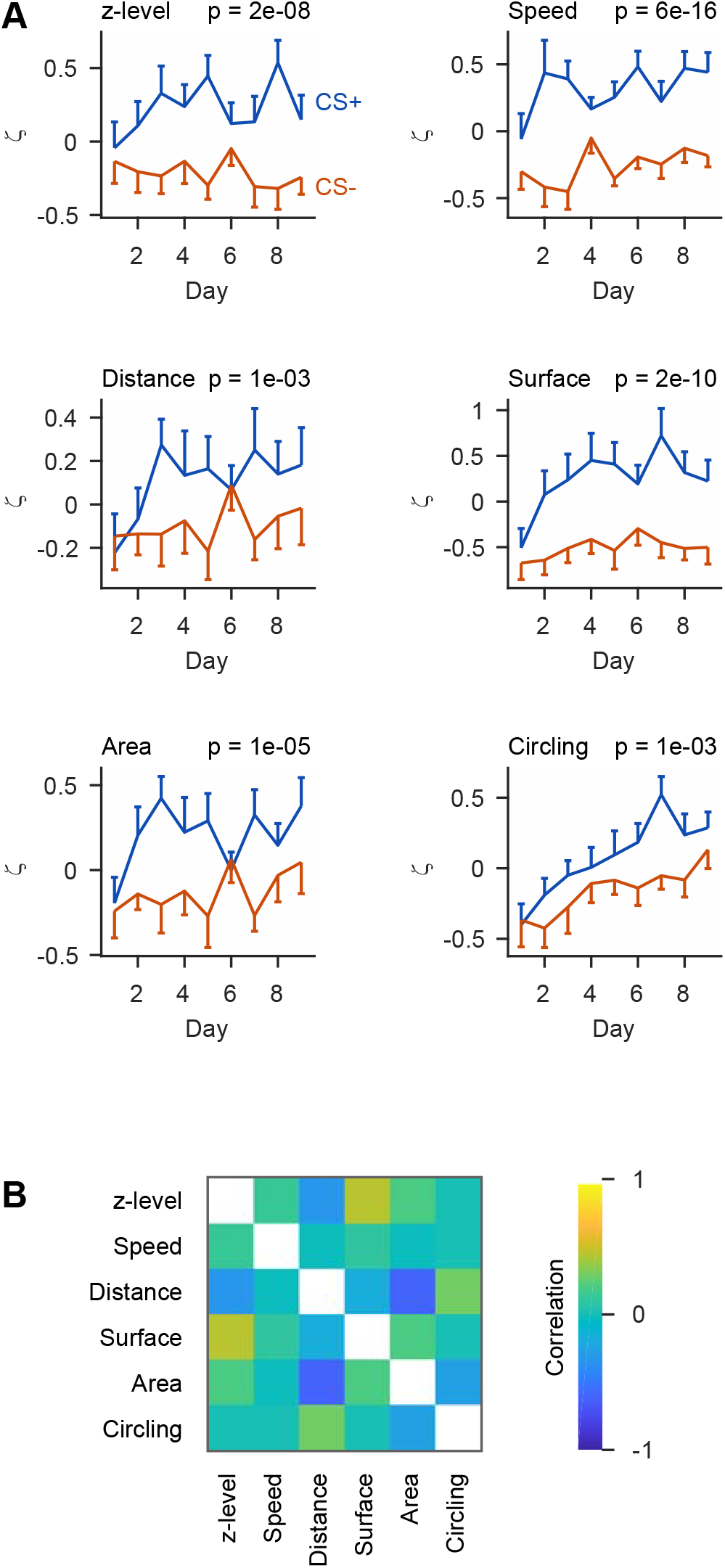
Analysis of discrimination learning. (A) ζ scores for the six behavioral parameters as a function of time, binned per day and averaged over fish (n = 12 fish). P-values show the probability that ζ scores differ between CS+ and CS- trials (Wilcoxon rank-sum test comparing all individual trials). (B) Mean correlation between time series of ζ scores for different behavioral parameters (n = 62 fish from experiments 1 – 3). Correlations were calculated on a trial-by-trial basis for each fish and averaged over fish.

**Fig. 4.**
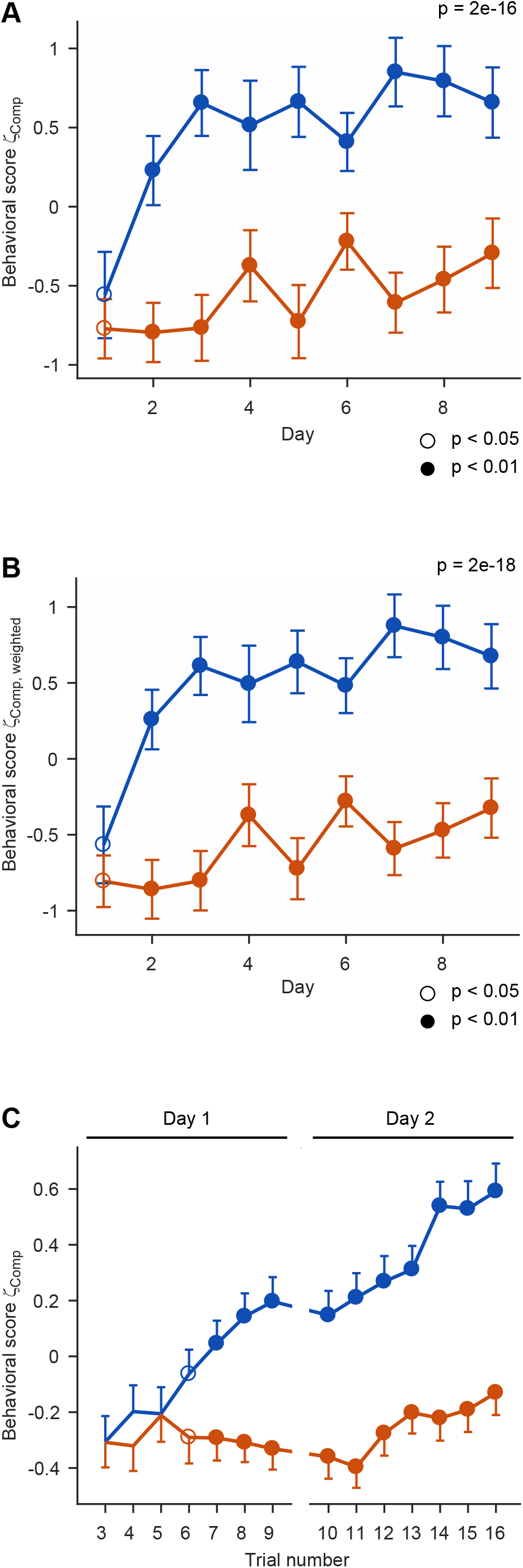
Discrimination learning curve. (A) Mean composite behavioral scores ζ_comp_ for responses to the CS+ and CS- as a function of training days (experiment 1). Data points for each day show the median of nine trials, averaged over n = 12 fish. Error bars show s.e.m. over fish. P-value gives the probability that ζ scores differ between CS+ and CS- trials (Wilcoxon rank-sum test comparing all individual trials, pooled over all days). Circles depict significant differences between behavioral responses to the CS+ and CS- on each day (Wilcoxon rank-sum test comparing all individual trials on a given day). Open circles: 0.01 ≤ p < 0.05; Filled circles: p < 0.01. (B) Same as above for the combined behavioral score ζ_comp,weighted_, which weights individual ζ scores by their auto/cross-covariances. (C) Quantification of behavioral responses to the CS+ and CS- in successive trials during the first 2 days of training, averaged over all fish in experiments 1 – 3 (n = 62 fish; mean ± s.e.m.). The first two trials and the last two trials were omitted because individual trial series of ζ scores were median-filtered with window size five to minimize the potential impact of outliers. Open and filled circles depict significant differences between behavioral responses to the CS+ and CS- as in (A).

The detection of learned behaviors may be further optimized by taking the covariances between behavioral parameters into account. To test this hypothesis we weighted each parameter by the ratio of its auto-covariance to the sum of its auto- and cross-covariances (Stouffer et al. 1949). Covariances were determined using all fish in experiments 1 – 3 (n = 62). The resulting combined score ζ_comp,weighted_ is thus corrected for redundant information in individual parameters. This approach further enhanced the separation of behavioral scores for the CS+ and CS- (Fig. 4B). However, the enhancement was minor, consistent with the modest cross-correlation between behavioral response components (Fig. 3B). The unweighted ζ_comp_ therefore offers the opportunity to obtain a highly informative combined behavioral score without the need to determine a behavioral covariance matrix, which is labor-intensive. As this approach is likely to be chosen in future applications, further analyses used the non-weighted measure ζ_comp_.

Odor discrimination at the end of training was quantified by the difference d between ζ_comp_ for the CS+ and CS-, averaged over the last 12 trials. After nine days of training, d was significantly different from zero (d = 0.98 ± 0.28; mean ± s.e.m.; n = 12 fish; p = 0.005; Wilcoxon signed rank test), confirming that fish discriminated between the CS+ and CS-.

Behavioral responses to the CS+ and CS- deviated already during the first day of training and approached saturation after approximately three days (Fig. 4A). Differences between behavioral scores for the CS+ and the CS- were statistically significant already on day 2, and the discrimination score d was statistically significant already when all data after day 2 were omitted (d = 0.66 ± 0.26; mean ± s.e.m.; n = 12 fish; p = 0.03; Wilcoxon signed rank test). Hence, fish learned to discriminate between odors after few training trials.

### 3.3. Experiments 2 and 3: shorter discrimination training

In order to corroborate these results we trained 36 additional fish on the discrimination of Ala and Trp for 27 – 36 trials (experiment 2). Again, fish showed significantly different behavioral responses to the CS+ and CS- at the end of training, with a discrimination score similar to that observed in experiment 1 (d = 1.14 ± 0.17; mean ± s.e.m.; n = 36 fish; p = 3e-7, Wilcoxon signed rank test; Fig. 5).

**Fig. 5.**
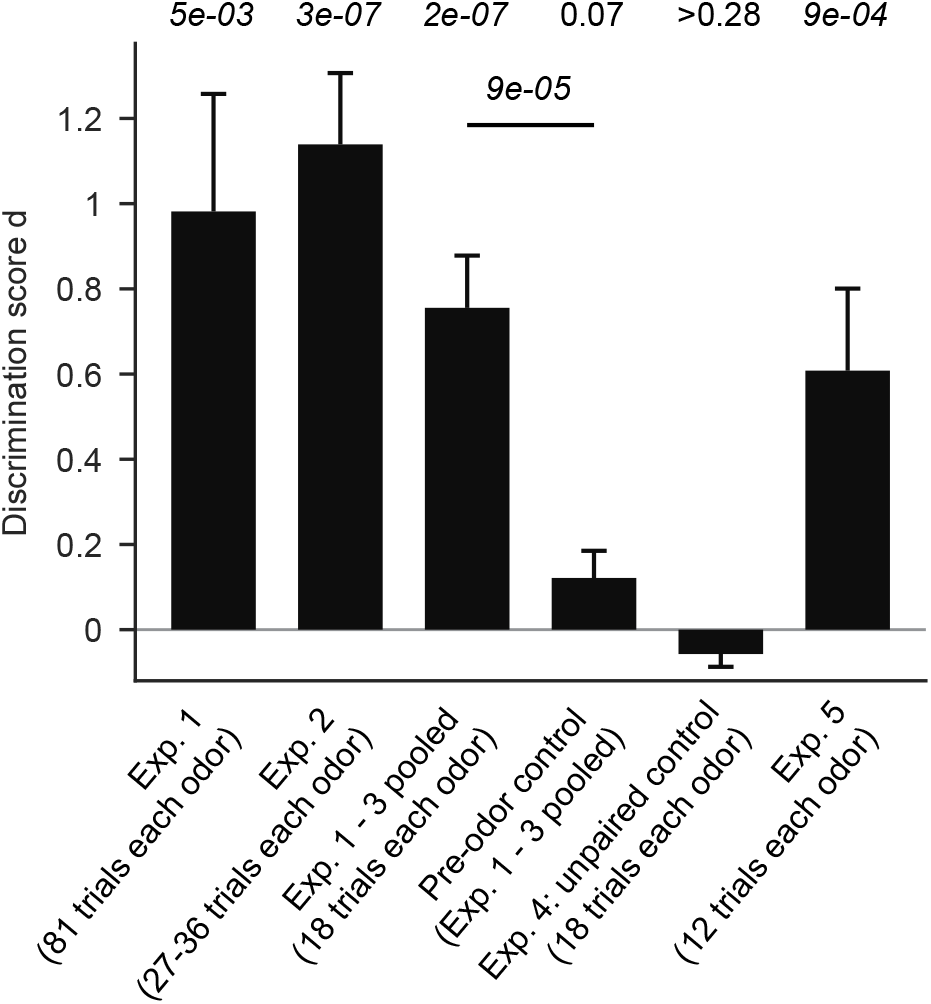
Quantification of discrimination behavior. Bars show the discrimination score d (mean ± s.e.m.) obtained in different experiments. P-values above show the probability that the mean discrimination score is significantly different from zero (Wilcoxon signed rank test). Horizontal line shows statistical comparison between the discrimination score d obtained in pooled experiments 1 – 3 (first 18 trials only) and the d value of the corresponding pre-odor control (Wilcoxon rank-sum test).

In addition, another 14 fish were trained on the discrimination of Ala and Trp for a total of 18 trials each (experiment 3) and showed significant discrimination behavior at the end of training (d = 0.51 ± 0.15; mean ± s.e.m.; n = 14 fish; p = 0.002, Wilcoxon signed rank test). These results were pooled with the first 18 CS+ and 18 CS- trials from experiments 1 and 2 to obtain a large sample (n = 62 fish). The mean discrimination score in this sample was (d = 0.76 ± 0.12; mean ± s.e.m.; p = 2e-7, Wilcoxon signed rank test; Fig. 5). The large sample size was exploited to analyze the initial phase of the learning curve at single-trial resolution. In order to minimize the impact of outliers, trial series of ζ_comp_ scores were median-filtered using a window size of five trials. The mean behavioral scores ζ_comp_ of responses to the CS+ and CS- gradually diverged and became statistically different at trial six (median-filtered data from trials 4 – 8; Fig. 4C). Hence, significant discrimination behavior emerged already during the first day of training.

### 3.4. Experiment 4: unpaired control

Differential responses to the CS+ and CS- in experiments 1 and 2 cannot be explained by innate odor preferences because amino acid stimuli were balanced across fish, implying that the observed behavior reflects associative learning. To confirm this conclusion we performed two controls. First, we analyzed behavior in the absence of a conditioned stimulus by re-analyzing swimming trajectories from experiments 1 – 3 using the time window 60 s to 30 s before odor application as the response time window. The baseline and reference time windows were shifted accordingly. The resulting value of d was not significantly different from zero (d = 0.12 ± 0.07; mean ± s.e.m.; n = 62 fish; p = 0.07, Wilcoxon signed rank test) and significantly different from the discrimination score of stimulus-induced behavior in the same fish (p = 9e-5; Fig. 5). Hence, the observed discrimination behavior cannot be explained by a bias in the training or analysis procedure.

Second, we exposed an additional 16 fish to 18 presentations of Ala and 18 presentations of Trp as before (9 trials each per day; two days total). However, food was delivered 15 min after application of both odors with a probability of 50%. Hence, fish were exposed to the same number of odor stimuli and food deliveries as before but odors did not predict food. We then randomly assigned Ala or Trp as CS+ and CS- in different fish in a balanced fashion and analyzed behavioral responses as before. The procedure was repeated 10 times for different random stimulus assignments. No significant discrimination score was obtained in any of these assignments (p > 0.28 in all cases; mean ± s.d.: p = 0.64 ± 0.22; range, 0.28 – 0.87) and the discrimination score remained close to zero (d = –0.06 ± 0.10; mean ± s.d.; Fig. 5). These results support the conclusion that the differential behavioral responses in experiments 1 – 3 reflect associative learning.

### 3.5. Experiment 5: conditioning can overcome innate preferences

We further examined whether the learned association between an odor stimulus and a food reward can overcome innate odor responses. A previous study reported that naïve zebrafish are attracted to Ala but repelled by Cys (Vitebsky et al. 2005). We tested whether the aversive response to Cys can be overcome when zebrafish are trained in the odor discrimination paradigm with Cys as CS+ and Ala as CS-. Each fish received only 12 trials with each stimulus, distributed over 3 – 5 days. Nevertheless, training resulted in the emergence of positive behavioral responses to Cys and odor discrimination was statistically significant (d = 0.61; n = 14 fish; p = 9e-4; Fig. 5). Hence, associative conditioning can establish appetitive behavioral responses to aversive odors.

## 4. Discussion

We developed efficient methods to analyze olfactory discrimination learning in adult zebrafish. Fish learned to respond selectively to one of two odor stimuli with anticipatory appetitive behavior. Training and analysis procedures are automated and can be performed in a standard laboratory environment. This paradigm is well-suited to analyze odor discrimination behavior and the underlying neuronal mechanisms in zebrafish.

### 4.1. Odor discrimination paradigm

Olfactory conditioning resulted in complex modifications of behavioral responses to odors. Learned responses to the CS+ included an increase in swimming velocity, an approach of the reward zone, and sampling of the water surface, reminiscent of the behavior that fish display in their home tanks when caretakers approach for feeding. Hence, fish appeared to learn the association between an olfactory cue and a familiar set of feeding-related behaviors. We therefore assume that our training procedure resulted primarily in classical conditioning.

Previous procedures for olfactory conditioning of adult zebrafish are difficult to implement in a standard laboratory setting because they require enormous amounts of water (Braubach et al. 2009) or very large tanks (Miklavc and Valentinčič 2012). Moreover, training and analysis were time-consuming because procedures are not automated. Importantly, previous procedures did not involve differential conditioning to two odors (CS+ and CS-), with few exceptions Miklavc and Valentinčič 2012). We therefore developed a procedure that is fully automated, requires only standard resources, and includes discrimination training. We expect that this procedure will be valuable to analyze the neural basis of discrimination learning.

A recent study by Doyle and colleagues described automated procedures for auditory and visual conditioning of adult zebrafish that also rely on the conditioning of feeding behavior (Doyle et al. 2017). The tasks follow a similar rationale as ours but exhibit differences in experimental procedures. First, we did not maintain fish in the fish facility during training, mainly to facilitate odor application and to avoid contamination of the circulating water with odors. Second, we trained fish individually rather than in groups. Training one fish per tank decreases throughput but facilitates quantitative analyses of multiple behavioral components and enables comparisons between individuals. Third, we trained fish to discriminate between two sensory cues (CS+ and CS-), rather than to associate a single cue with a behavioral output, because differential conditioning is desired to analyze the neural basis of sensory discrimination. Despite the differences in task design and sensory modalities, fish rapidly learned associations between the CS and US in both studies (Doyle et al. 2017). An appetitive paradigm for operant visual discrimination learning, in contrast, required more extensive training (Mueller and Neuhauss 2012), possibly because fish needed to learn a novel behavioral sequence to collect rewards. Hence, classical conditioning of feeding behavior appears to be an effective strategy for the design of appetitive learning tasks in zebrafish.

Based on observations of behaving fish we hypothesized that the conditioned behavior consists of multiple behavioral components, and that the contribution of different behavioral components varies between trials and individuals. Consistent with this observation, the combined analysis of multiple behavioral components improved the detection of learned behavioral responses. Hence, the overt behavior was complex even though the conditioning paradigm is conceptually simple.

### 4.2. Olfactory discrimination learning

Zebrafish developed differential behavioral responses to related amino acid odors when and only when they differed in their prediction of food. Hence, fish discriminated between odors and learned specific associations between odors and behaviors, consistent with previous findings (Braubach et al. 2009; Miklavc and Valentinčič 2012). Appetitive olfactory conditioning could be achieved even using an odor that is innately aversive. Hence, olfactory conditioning is a robust phenomenon that can overcome innate odor preferences. This approach may be exploited to dissect neural pathways that mediate innate and learned behavioral responses to odors.

The 30 s delay between the onset of the CS and US provided the opportunity to quantify behavioral responses in each trial. Continuous learning curves could therefore be acquired without the need to introduce separate probe trials, which may result in extinction. As the acquisition of memory can be sensitive to the precise temporal relationship between the CS and the US, experimental conditions may be further improved by optimizing the delay between odor and food application. However, we did not attempt this because the chosen protocol already resulted in robust conditioning.

Statistically different behavioral responses to the CS+ and CS- were detected already after few training trials. Zebrafish can therefore establish specific olfactory memories based on a small amount of experience. Odor discrimination tasks that are widely used in rodents, in contrast, often require hundreds of training trials to reach asymptotic performance (Abraham et al. 2004; Bodyak and Slotnick 1999; Rinberg et al. 2006). The reason for this difference in training requirement remains unclear. One possibility is that the tasks for rodents include unnatural behavioral components such as nose pokes whereas the task for fish is based on the conditioning of familiar food search behavior.

The ability of animals to discriminate odors is usually assessed by a behavioral readout that is based on odor discrimination learning. However, because learning itself modifies odor representations (Abraham et al. 2014; Chapuis and Wilson 2011; Chu et al. 2016; Li et al. 2008; Yamada et al. 2017), mechanisms of odor discrimination in naïve or nearly naïve animals remain difficult to analyze. A rapid conditioning procedure may open new opportunities to address this issue. We anticipate that the combination of behavioral and physiological analyses in zebrafish will provide new insights into the neuronal basis of olfactory processing and memory.

## Acknowledgements

We thank G. Jacobson for helpful input and support and the Friedrich lab for stimulating discussions. This work was supported by the Novartis Research Foundation, by the Deutsche Forschungsgemeinschaft (DFG; FR 1667/2-2), by the Swiss National Science Foundation (SNF; 31003A_135196 and 310030B_1528331), and by the European Research Council (ERC) under the European Union’s Horizon 2020 research and innovation programme (grant agreement No 742576).

